# A unity of opposites in between Nrf1- and Nrf2-mediated responses to the endoplasmic reticulum stressor tunicamycin

**DOI:** 10.1101/655159

**Authors:** Yu-ping Zhu, Shaofan Hu, Xufang Ru, Ze Zheng, Zhuo Fan, Lu Qiu, Yiguo Zhang

## Abstract

The water-soluble Nrf2 is accepted as a master regulator of antioxidant responses to cellular stress, it was also identified as a direct target of the endoplasmic reticulum (ER)-anchored PERK. However, the membrane-bound Nrf1 response to ER stress remains elusive. Herein, we report a unity of opposites in both Nrf1- and Nrf2-coordinated responses to the ER stressor tunicamycin (TU). The TU-inducible transcription of Nrf1 and Nrf2, as well as GCLM and HO-1, was accompanied by activation of ER stress signaling networks. The unfolded protein response (UPR) mediated by ATF6, IRE1 and PERK was significantly suppressed by Nrf1α-specific knockout, but hyper-expression of Nrf2, GCLM and HO-1 was retained in *Nrf1α*^*−/−*^ cells. By contrast, *Nrf2*^*−/−ΔTA*^ cells with a genomic deletion of its transactivation domain resulted in significant decreases of GCLM, HO-1 and Nrf1; this was accompanied by partial decreases of IRE1 and ATF6, but not PERK, along with an obvious increase of ATF4. Notably, Nrf1 glycosylation and its *trans*-activity to mediate transcriptional expression of 26S proteasomal subunits were repressed by TU. This inhibitory effect was enhanced by *Nrf1α*^*−/−*^ and *Nrf2*^*−/−ΔTA*^, but not by a constitutive activator *caNrf2*^*ΔN*^ (that increased abundances of non-glycosylated and processed Nrf1). Furthermore, *caNrf2*^*ΔN*^ also enhanced induction of PERK and IRE1 by TU, but reduced expression of ATF4 and HO-1. Such distinct roles of Nrf1 and Nrf2 are unified to maintain cell homeostasis by a series of coordinated ER-to-nuclear signaling responses to TU. Overall, Nrf1α acts in a cell-autonomous manner to determine transcription of most of UPR-target genes, albeit Nrf2 is also partially involved in this process.

## Introduction

As a highly-evolved organelle of eukaryotic cells, endoplasmic reticulum (ER) is of crucial importance to be involved in biosynthesis of secretory and membrane proteins, as well as lipids including cholesterol, proper folding of proteins and specific post-translational modifications of those proteins sorted out of the ER to their destination organelles. The ability of the ER to orchestrate intracellular proteins and lipids is also severely challenged by a vast variety of physio-pathological stresses and environmental stimuli [1]. Consequently, disruption of such function of ER leads to accumulation of a large amount of unfolded and/or misfolded proteins in the lumen of this organelle, destroying the original homeostasis of the cell; this is known as ER stress to activate the unfolded protein response (UPR) [2]. In metazoans, the UPR is mediated by three major axes of signaling pathways from: i) PERK (PRKR-like ER kinase)-eIF2α (eukaryotic translation initiation factor 2α) to ATF4 and Chop; ii) IRE1 (inositol-requiring protein 1) to XBP1 (X-box binding protein 1); and iii) ATF6 [3]. In early stages of UPR, those unfolded proteins are allowed for binding to an ER luminal-resident chaperone BIP (immunoglobulin heavy chain-binding protein homolog, also called glucose regulated protein 78, i.e. GRP78) [4]. This critical event results in the induction of those ER-associated sensors PERK, IRE1 and ATF6, so that their relevant UPR signaling pathways are subsequently activated [5] (as illustrated in Figure 1).

**Figure 1.**
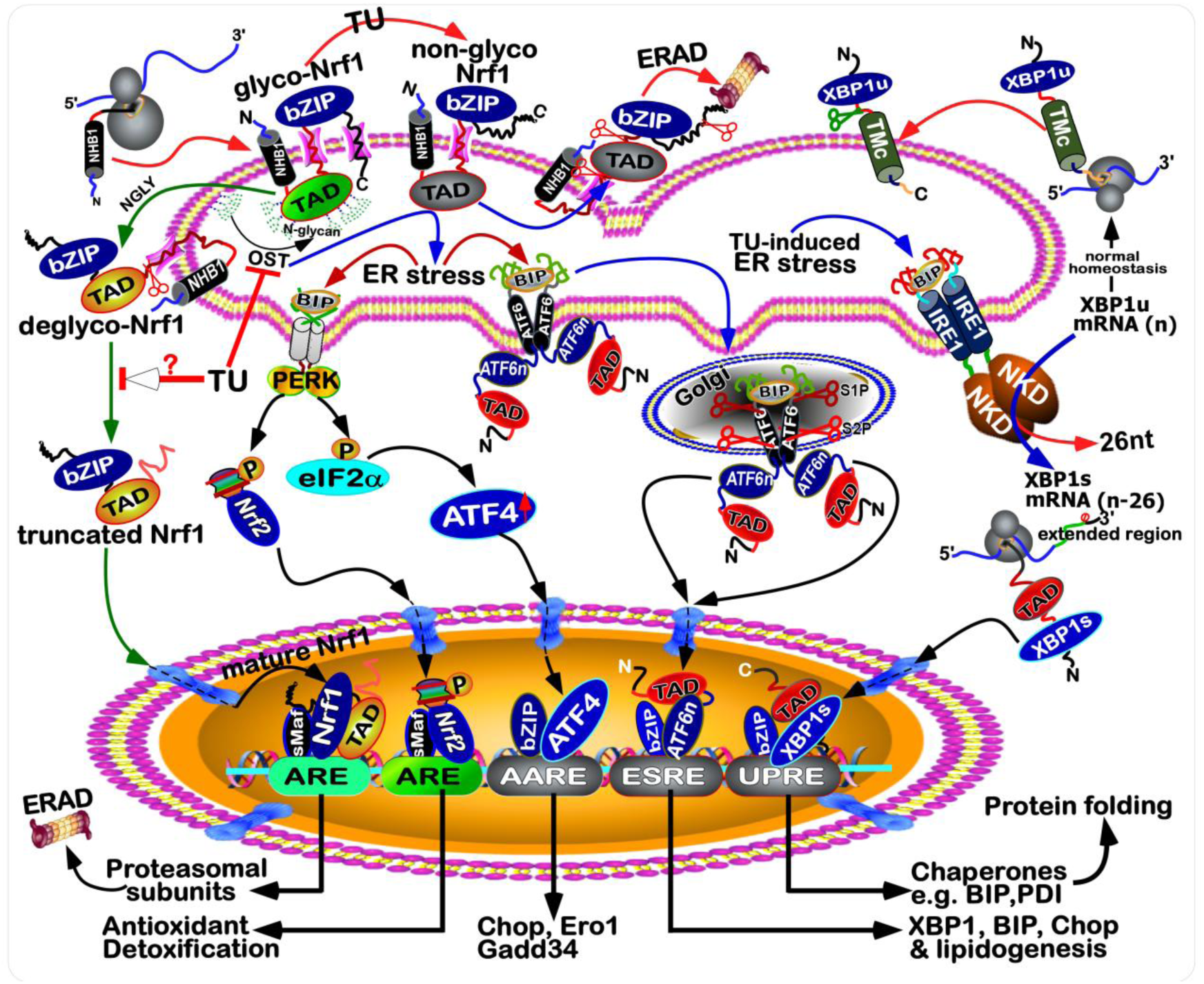
Inhibition of Nrf1 N-glycosylation by TU in the ER-to-nuclear signaling responses. A model is herein proposed to explain the inhibition of Nrf1 N-glycosylation by TU, as a classic stressor, in order to block its deglycosylation and proteolytic processing to yield a mature CNC-bZIP factor. The resulting non-glycosylated Nrf1 and others are subject to the ER-associated degradation (ERAD) pathway. If the non-glycosylated and unfolded proteins are accumulated so much as to stimulate ER stress. Thereby, inhibition of Nrf1 by TU is also accompanied by activation of canonic ER-to-nuclear response signaling mediated by PERK, IRE1 and ATF6, as reviewed by the authors [1-4]. Of note, PERK was also identified as a direct upstream kinase of Nrf2, as well as elF2α [10], albeit this CNC-bZIP protein has been well-characterized as a master antioxidant transcription factor to regulate ARE-driven genes. In this response, ATF4 is selectively translated by the elF2α-control machinery to regulate target genes (including *Chop, Ero1, Gadd34*) by their amino acid response elements (AARE) in the promoter regions. To get rid of ER stress, ATF6 was activated by consecutive proteolytic processing of this bZIP protein by Site-1 and Site-2 proteases (i.e., S1P and S2P). The full-length mRNA of XBP is alternatively spliced to remove its 26 nucleotides at nearly 3′-end, such that the subsequent portion of its open reading frame is shifted to yield a longer inducible protein isoform called XBP-1s than its original protein of XBP-1u. Consequently, distinct subsets of cognate target genes containing either *cis-*regulatory ESRE (ER stress response element) or UPRE (unfolded protein response element) are regulated transcriptionally by ATF6n (i.e., its active N-terminal portion) and XBP-1s, respectively.

Upon activation of UPR, it is also accompanied by concomitant and secondary induction of a complex signaling networks, which include pro-survival mechanisms involving antioxidant responses, ER-associated degradation (ERAD) ER biogenesis and autophagy [6]. Notably, the potential role of UPR in maintaining redox homeostasis is attracting great interest of workers in different fields [7]. Clearly, it is generally accepted that redox responses are mediated by two important antioxidant transcription factors Nrf1 [i.e. nuclear factor erythroid 2 (NF-E2) p45-related factor 1, also called Nfe2l1] and Nrf2 [8]. These two members of the cap’n’collar (CNC) basic-region leucine zipper (bZIP) family can predominantly regulate transcription of antioxidant response element (ARE)-driven genes involved in detoxification and other cytoprotective adaptations. Besides, Nrf2 was also previously reported to be significantly up-regulated by amyloid beta-induced ER stress leading to UPR [9]. As one of the three UPR transducers, PERK had been identified to be a direct upstream kinase of Nrf2 [10], which is well characterized as a master antioxidant transcription factor to counterbalance the harmful effects of reactive oxygen species (ROS) on cells. Under ER stress, Nrf2 is phosphorylated by PERK, such that the phosphorylated Nrf2 dissociates from its negative regulator KEAP1 and then translocated the nucleus leading to transactivation of ARE-battery genes [11, 12]. In the meanwhile, eIF2α is also phosphorylated by PERK, thereby repressing general protein translation, but promoting selective protein translation of ATF4 [13-15]. Furtherly, the heterodimer consisting of ATF4 and Nrf2 binds to the stress-response element of heme oxidase-1 (HO-1), before inducing this gene expression [16]. Collectively, these demonstrate that Nrf2-mediated expression of ARE genes is activated in the UPR signaling to ER stress. However, whether (and how) the ER membrane-associated Nrf1-mediated response signaling is triggered to the accompaniment of UPR remains elusive, to date.

Although the ER-associated Nrf1 adapts a unique membrane-topology (as shown in Figure 1), at least in part of which is similar to the consensus transmembrane topology of PERK, IRE1 and ATF6. Thereby, it is postulated that the membrane-bound Nrf1 should, theoretically, has a strong capacity of being induced by ER stress, as described for the homologue of *Caenorhabditis elegans* Skn-1 [17-19]. Intriguingly, ectopic Nrf1 appeared to be *de facto* not activated by each of UPR signaling pathways, but conversely, activation of Nrf1 by *tert*-butylhydroquinone (tBHQ) or arsenic to up-regulate ARE-driven genes responsible for antioxidant, detoxification and cytoprotection was repressed by classic ER stressors, such as tunicamycin (TU), thapsigargin (TG) and brefeldin A (BFA) [20, 21]. As such, the electrophoretic mobility of Nrf1 and its subcellular redistribution were altered by TU and BFA, but not TG [21]. This suggests that Nrf1 is modified in an ER stress-stimulated post-translational fashion (i.e. N-glycosylation, deglycosylation, ubiquitination, and proteolysis). By sharp contrast, the water-soluble Nrf2 was identified as a direct target of PERK triggered by TU, insofar as to activate antioxidant and detoxification responses [10, 12].

It is important to note a unique UPR-independent mechanism whereby the transactivation activity of Nrf1, rather than Nrf2, to up-regulate transcription of all those genes encoding 26S proteasomal (*PSM*) subunits is induced by low concentrations of its inhibitors [22-24]. Such being the case, proteasomal inhibition can also result in accumulation of oxidative ubiquitinated proteins, thereby triggering ER stress [24-26]. In fact, ER stress-inducible UPR signaling pathways were endogenously activated by loss of mouse Nrf1 function in the homozygous (*Nrf1*^*−/−*^) hepatocytes [27]. Contrarily, both similar ER stress responses and steatosis were enhanced by half loss of Nrf1 in the heterozygous (*Nrf1*^*+/−*^) livers, when compared with wild-type (*Nrf1*^*+/+*^) livers, in response to 26S proteasomal inhibition [27]. Such concurrence of severe oxidative stress with UPR in mouse *Nrf1*^*−/−*^ livers results from down-regulation of antioxidant, detoxification and proteasomal genes. In the proteasomal compensatory response to limited extents of proteasome inhibitors [28], Nrf1 is subjected to the multistep processing of Nrf1 and its nuclear translocation [29], albeit the inhibition of proteasome-mediated ERAD is also one of the causes of ER stress [24, 26, 30]. These facts demonstrate that Nrf1 possesses an essential cytoprotective role against development of hepatosteatosis by maintaining the ER homeostasis, but the underlying mechanism remains unknown.

To unveil the mystery players of Nrf1 and Nrf2 in UPR, it is necessary to gain insights into distinct roles of both CNC-bZIP factors in the ER response to TU. Herein, we report a unity of opposite roles of Nrf1 and Nrf2 in the cellular responses to treatment of TU in different genotypic cell lines. TU-induced UPR signaling by ATF6, IRE1 and PERK was significantly suppressed by Nrf1α-specific knockout, but Nrf2, GCLM and HO-1 was highly expressed in *Nrf1α*^*−/−*^ cells. By contrast, *Nrf2*^*−/−ΔTA*^ cells with a genomic deletion of its transactivation domain resulted in significant decreases of GCLM, HO-1 and Nrf1. This was also accompanied by partial decreases of IRE1 and ATF6, but not PERK, along with an obvious increase of ATF4. Notably, glycosylation of Nrf1 and its ability to transactivate expression of 26S proteasomal subunits were markedly repressed by TU. This inhibitory effect was enhanced by *Nrf1α*^*−/−*^ and *Nrf2*^*−/−ΔTA*^, but not by a constitutive activator *caNrf2*^*ΔN*^ because of increased abundances of non-glycosylated and processed Nrf1. In addition, *caNrf2*^*ΔN*^ enhanced induction of PERK and IRE1 by TU, but reduced ATF4 and HO-1. Collectively, such distinctive roles of Nrf1 and Nrf2 in the ER-to-nuclear signaling responses to TU are unified for maintaining cell homeostasis. Overall, our results demonstrate that Nrf1α acts as a dominant player in a cell-autonomous manner to regulate most of UPR gene expression, while Nrf2 is partially involved in this process by IRE1, at least in this experimental setting.

## Results

### Nrf1α and Nrf2 contribute to differential expression of responsive genes to ER stress in four different cell lines

To explore distinct functions of Nrf1 and Nrf2 in the putative ER stress response, herein we employed four different genotypic cell lines, which had been established by gene-editing with the presence or absence of Nrf1α and Nrf2 [31]. Consistently, these selected cell lines were re-identified by Western blotting of Nrf1 and Nrf2before RNA sequencing. As shown in Figure 2A, wild-type HepG2 cell line (*Nrf1/2*^*+/+*^) served as a control for the expression of Nrf1 and Nrf2. By contrast, *Nrf1α*^*−/−*^ cell line exhibited a clear disappearance of the intact full-length Nrf1α/TCF11 and its derivatives (which are embodied by glycoprotein-A, deglycoprotein-B and N-terminally-truncated C-/D-forms of between ∼140- and 120-kDa; their detailed identifications had been described in our previous work [24, 32]). However, expression of shorter Nrf1-truncated E-form appeared to be enhanced in *Nrf1α*^*−/−*^ cell with unaltered F-form, when compared with *Nrf1/2*^*+/+*^ cells (Figure 2A, *upper panel*). This implies a potential molecular compensatory mechanism, because E- and F-isoforms of Nrf1 may be generated by translation of a shorter-length open reading frame of mRNA resulting from the alternative first exon (i.e. Nrf1^ΔN^) [8, 24], in addition to proteolytic processing of longer Nrf1 isoforms. Intriguingly, almost no expression of the Nrf1-truncated E-isoform was determined in both cell lines of *Nrf2*^*−/−ΔTA*^ and *caNrf2*^*ΔN*^, but they gave a modest decrease in abundance of Nrf1 F-form. Similarly, abundances of Nrf1β bands close to 70-kDa appeared to be unaffected by knockout of Nrf1α, but was significantly augmented by *Nrf2*^*−/−ΔTA*^ and rather reduced by *caNrf2*^*ΔN*^ (Figure 2A, *upper panel*). These data suggest a potential effect of Nrf2 on alternative translation of either Nrf1^ΔN^ or Nrf1β, but another possible role of Nrf2 in alternative transcription of Nrf1 cannot also be ruled out, albeit detailed mechanism(s) remains unknown.

**Figure 2.**
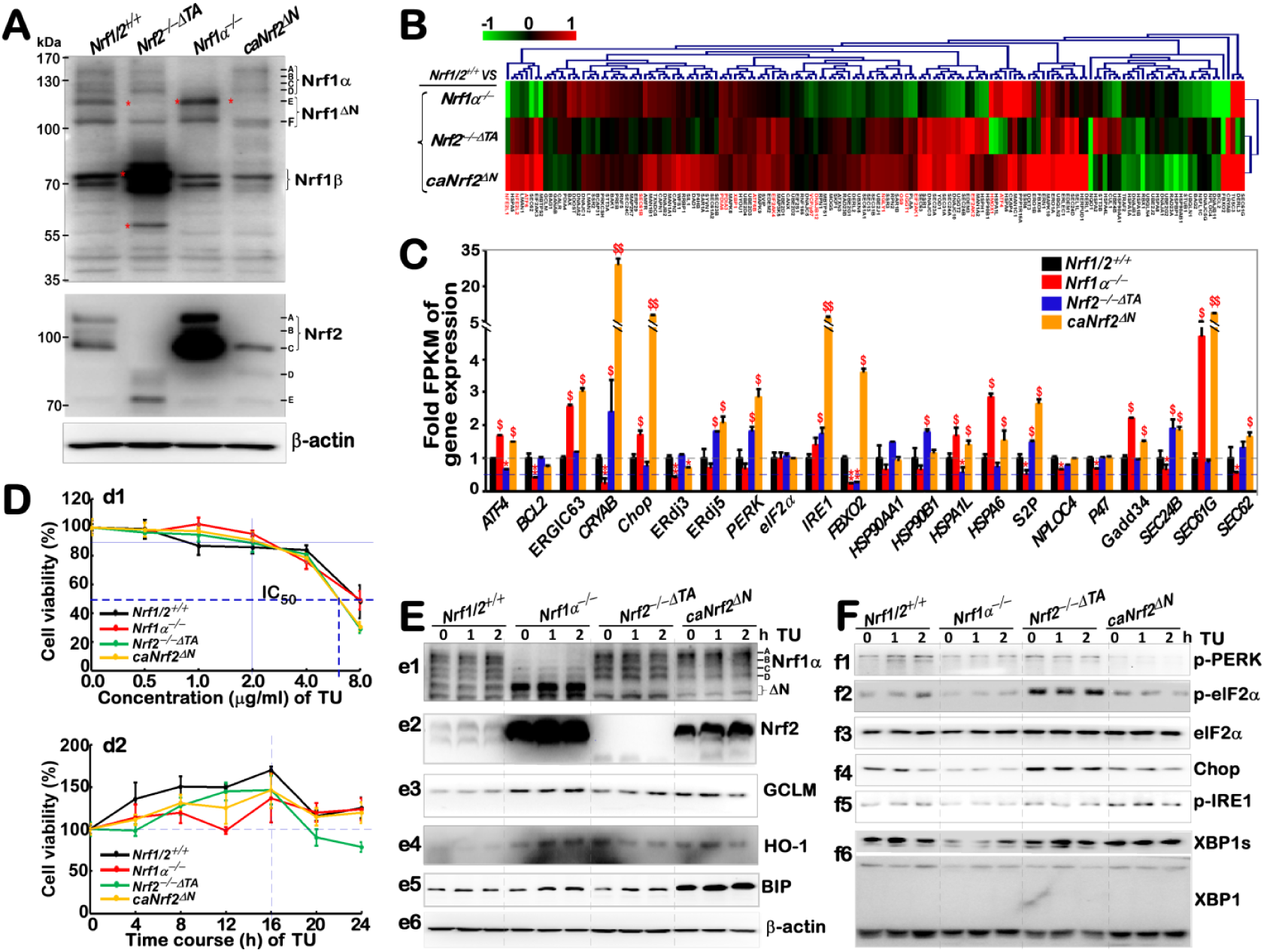
Distinct contributions of Nrf1α and Nrf2 to differential expression of ER stress-related genes. (A) Distinct protein levels of Nrf1 and Nrf2 in different genotypic cell lines *Nrf1/2*^*+/+*^, *Nrf1α*^*−/−*^, *Nrf2* ^*−/−δTA*^ and *caNrf2*^*δN*^ were determined by Western blotting with specific antibodies. (B) A heatmap was made by the Log2-based RPKM values, representing differential expression profiles of ER stress-related genes by comparison to those obtained from *Nrf1*^*+/+*^ cells. Different changes in the basal expression of these genes are shown to distinct degrees of colors. (C) The expression of ER stress responsive genes was also evaluated by relative RPKM values. Significant statistical decreases were indicated by \**p*<0.01 or \*\**p*<0.001, whereas significant increases were represented by *$*, p<0.01 or *$$, p*<0.001. (D) The viability of these cell lines that had been treated with different concentrations of TU for 24 h (*d1*), or treated with2 μg/ml of TU for different lengths of time (*d2*), were determined by an MTT-based assay. (E) The protein levels of antioxidant genes (i.e. *e1 to e6*) and (F) the ER stress-related genes (i.e. *f1 to f6*) in the experimental cells, that had been treated with 2 μg/ml of TU for a short time (from 0 to 2 h), were examined by Western blotting with indicated different antibodies.

By contrast with *Nrf1/2*^*+/+*^ cells, *Nrf1α*^*−/−*^ cells gave rise to a dramatic increase in the expression of Nrf2 between ∼100- and 110-kDa (Figure 2A, *middle panel*); such a demonstrating effect of Nrf1 on the expression of Nrf2 was also described previously [31]. Nonetheless, similar longer Nrf2 between ∼100- and 110-kDa were completely abolished in *Nrf2*^*−/−ΔTA*^ cells but replaced by additional smaller molecular weight polypeptides with a genomic deletion of its transactivation domain. Notably, the intact full-length Nrf2 of ∼110-kDa was also totally abolished by *caNrf2*^*ΔN*^, but it retained a major short Nrf2 of ∼100-kDa, together with a few of minor small polypeptides (Figure 2A, *middle panel*). These suggest a possible proteolytic processing of Nrf2 within its N-terminal Neh2 domain (which is highly conserved with the Neh2L region immediately adjacent to the N-terminal domain of Nrf1).

Subsequently, the fidelity of total RNAs purified from the above-identified four cell lines was rigidly confirmed to be available for RNA-sequencing. A heatmap of the sequencing data revealed 157 of differentially expressed genes clustered responsibly for ER stress in *Nrf1α*^*−/−*^, *Nrf2*^*−/−ΔTA*^ *or caNrf2*^*ΔN*^ by comparison with *Nrf1/2*^*+/+*^ cells (Figure 2B). Amongst them, relative highly expressed genes were differed in these four cell lines as shown graphically (Figure 2C). Further analysis of these data, taken together with the aforementioned alternations in abundances of Nrf1 and Nrf2, suggests that basal expression of 7 genes encoding ATF4, Chop, Gadd34, ERGIC63, HSPA1L, HSPA6 and Sec61γ could be regulated predominantly by Nrf2, because their mRNA levels were significantly increased in *Nrf1α*^*−/−*^ and *caNrf2*^*ΔN*^ cells, but obviously diminished or abolished in *Nrf2*^*−/−ΔTA*^ cells (Figure 2C). By contrast, Nrf1α/TCF11 may be primarily involved in regulating basal expression of another 9 genes encoding BCL2, CRYAB, ERdj3, PERK, FBXO2, S2P, NPLOC4, Sec24β, and Sec62. This is due to the fact that their basal mRNA abundances were markedly repressed in *Nrf1α*^*−/−*^ cells, albeit with high expression of Nrf2. Conversely, a few of these genes regulated by Nrf1α might also be inhibited by Nrf2, because their mRNA expression levels were strikingly recovered by *Nrf2*^*−/−ΔTA*^ (or *caNrf2*^*ΔN*^ with a constitutive deletion of cytoplasmic Keap1-binding Neh2 domain of Nrf2, with an unidentified nuclear function of this domain). Notably, IRE1 could be co-regulated by both Nrf1α and Nrf2, because its mRNA expression levels were unaffected by knockout of Nrf1 or Nrf2, but significantly elevated by *caNrf2*^*ΔN*^, implying a possible release of inhibition by Neh2.

### Distinct effects of Nrf1α and Nrf2 in protein expression of responsive genes to early TU-induced ER stress

To determine effects of Nrf1 and Nrf2 on differential expression of putative genes in response to the ER stressor TU, we performed Western blotting to examine changes in the protein levels of early TU-responsive genes expressed, for a short period of time, in the aforementioned four cell lines with the presence or absence of Nrf1α and Nrf2. Firstly, cytotoxity of TU was evaluated to provide an optimal concentration of this chemical that was treated for an optimal time course (Figure 2D). A half of maximal inhibitory concentration (IC_50_) of TU treated in *Nrf2*^*−/−ΔTA*^ or *caNrf2*^*ΔN*^ cells was ∼6.5 μg/ml, while another IC_50_ of TU treatment of *Nrf1α*^*−/−*^ or *Nrf1/2*^*+/+*^ cells was close to 8.0 μg/ml (Figure 2D1). Of note, 2.0 μg/ml of TU, with almost no obvious cytotoxity for 24 h (Figure 2D2), was selected for the use of our subsequent experiments.

Western blotting revealed a modest increase in abundances of longer inactive Nrf1 isoform-A/B was examined following 2-h TU-treatment of *Nrf1/2*^*+/+*^ or *caNrf2*^*ΔN*^, but not *Nrf2*^*−/−ΔTA*^ cells (Figure 2E1); this appeared consistent with our previous work [24, 33]. As such, two shorter active isoforms-C/D of Nrf1 was enhanced by TU treatment of *Nrf2*^*−/−ΔTA*^ cells only. Notably, either Nrf2 or caNrf2^ΔN^ protein levels were promoted by TU treatment of *Nrf1α*^*−/−*^ or *caNrf2*^*ΔN*^ cells, respectively (Figure 2E2). Further examination unraveled that the protein expression of ARE-driven genes encoding GCLM and HO-1 (regulated Nrf1 and/or Nrf2) was significantly increased by TU treatment of *Nrf1α*^*−/−*^ and *caNrf2*^*ΔN*^ cells, when compared with *Nrf1/2*^*+/+*^ cells (Figure 2E3 and 2E4). Conversely, this antioxidant effect was partially reduced in TU-treated *Nrf2*^*−/−ΔTA*^ cells. Taken altogether, these data demonstrate that short-term stimulation of TU can also trigger induction of antioxidant gene response mediated by Nrf1 and Nrf2.

Furtherly, distinct time-dependant increases in the chaperone BIP (also called GRP78, as a landscape signature of classic TU-induced ER stress response) were determined following TU treatment of *caNrf2*^*ΔN*^, *Nrf1α*^*−/−*^, *Nrf2*^*−/−ΔTA*^ or *Nrf1/2*^*+/+*^ cells, when compared with their untreated controls (Figure 2E5). Of note, higher expression levels of basal and TU-stimulated BIP proteins was found in *caNrf2*^*ΔN*^ cells. By contrast, significant elevation of Nrf2 in *Nrf1α*^*−/−*^ cells only gave rise to a considerable level of BIP, but was only slightly suppressed by knockout of Nrf2 (in *Nrf2*^*−/−ΔTA*^ cells) to a similar level to that obtained from *Nrf1/2*^*+/+*^ cells. Collectively, these data indicate that both Nrf1 and Nrf2 are required for BIP expression in a short-term ER stress response to TU. Western blotting examination of the PERK-eIF2α/Nrf2 -Chop pathway revealed that both phosphorylated PERK and eIF2α proteins, along with total Chop were stimulated by TU treatment in *Nrf1/2*^*+/+*^ cells (Figure 2F1, 2F2 and 2F4). Interestingly, their expression was obviously attenuated by hyper-expression of Nrf2 in *Nrf1α*^*−/−*^ or *caNrf2*^*ΔN*^ cells, but also significantly recovered by *Nrf2*^*−/−ΔTA*^ to considerable high levels relative to those obtained from *Nrf1/2*^*+/+*^ cells; such alternations appears to be independent of stimulation by TU. No matter what it is, these observations demonstrate that Nrf1 and Nrf2 have exerted opposing roles for the PERK-eIF2α-Chop signaling response to a short-term stimulation of TU. Further examination of the IRE1-XBP1 pathway unraveled that basal and TU-stimulated expression of p-IRE and XBP1s appeared to be reduced in *Nrf1α*^*−/−*^ cells (Figure 2F5 and 2F6), but p-IRE1 expression was enhanced in *caNrf2*^*ΔN*^ cells, albeit no changes of both proteins were observed in *Nrf2*^*−/−ΔTA*^ by comparison with equivalents of *Nrf1/2*^*+/+*^ cells. Overall, these results indicate that Nrf1α, but not Nrf2, is required for expression of auto-phosphorylated IRE1 and its target product XBP1s.

### Nrf1α and Nrf2 mediate distinct transcriptional responses to the long-term TU-stimulated ER stress

The above experiments showed no changes in protein expression levels of some responsive genes at the early stages of TU-induced ER stress (Figure 2E,2F). Particularly, a few of them were *de facto* auto-activated in a TU-independent manner under the untreated homeostatic conditions. Thereby, the TU-treated time was further extended from 4 h to 24 h, in order to determine distinctions in between Nrf1- and Nrf2-mediated transcriptional responses to a long-term TU-stimulated ER stress. For this, four different genotypic cell lines had been treated with TU for distinct lengths of time from 0 to 24 h, before these genotypic mRNAs were subjected to real-time quantitative PCR (qRT-PCR) analysis. The results demonstrated that TU treatment of *Nrf1/2*^*+/+*^ cells triggered time-dependent increases in transcriptional expression of *Nrf1* and *Nrf2* (Figure 3A, 3B). By sharp contrast, both basal and TU-stimulated mRNA expression levels of *Nrf1* were substantially abolished by *Nrf1α*^*−/−*^ and also significantly diminished by *Nrf2*^*−/−ΔTA*^, but not by *caNrf2*^*ΔN*^ (Figure 3A). This further supports our previous notion that the *Nrf1* gene transcription is regulated positively by itself factor Nrf1α (and derivatives) together with Nrf2, as described by Qiu *et al* [31*]. Conversely, basal mRNA expression of Nrf2* was elevated in *Nrf1α*^*−/−*^ cells (with high abundances of Nrf2) to a 3.5-fold extent, as compared to the control value of *Nrf1/2*^*+/+*^ cells (Figure 3B). TU-treated *Nrf1α*^*−/−*^ cells only gave rise to a modest increase in expression of *Nrf2* mRNA following treatment for 8 h to 16 h, but subsequently, this increased expression was significantly suppressed to a lower level than its basal value. However, basal and TU-inducible mRNA expression levels of *Nrf2* were enhanced by *caNrf2*^*ΔN*^ with no decreases in its induction, but all completely abolished by *Nrf2*^*−/−ΔTA*^ (Figure 3B). These collective results indicate that transcription of *Nrf2* gene is regulated positively by itself Nrf2 factor, but also negatively (at least in part) by Nrf1 and/or its downstream factors.

**Figure 3.**
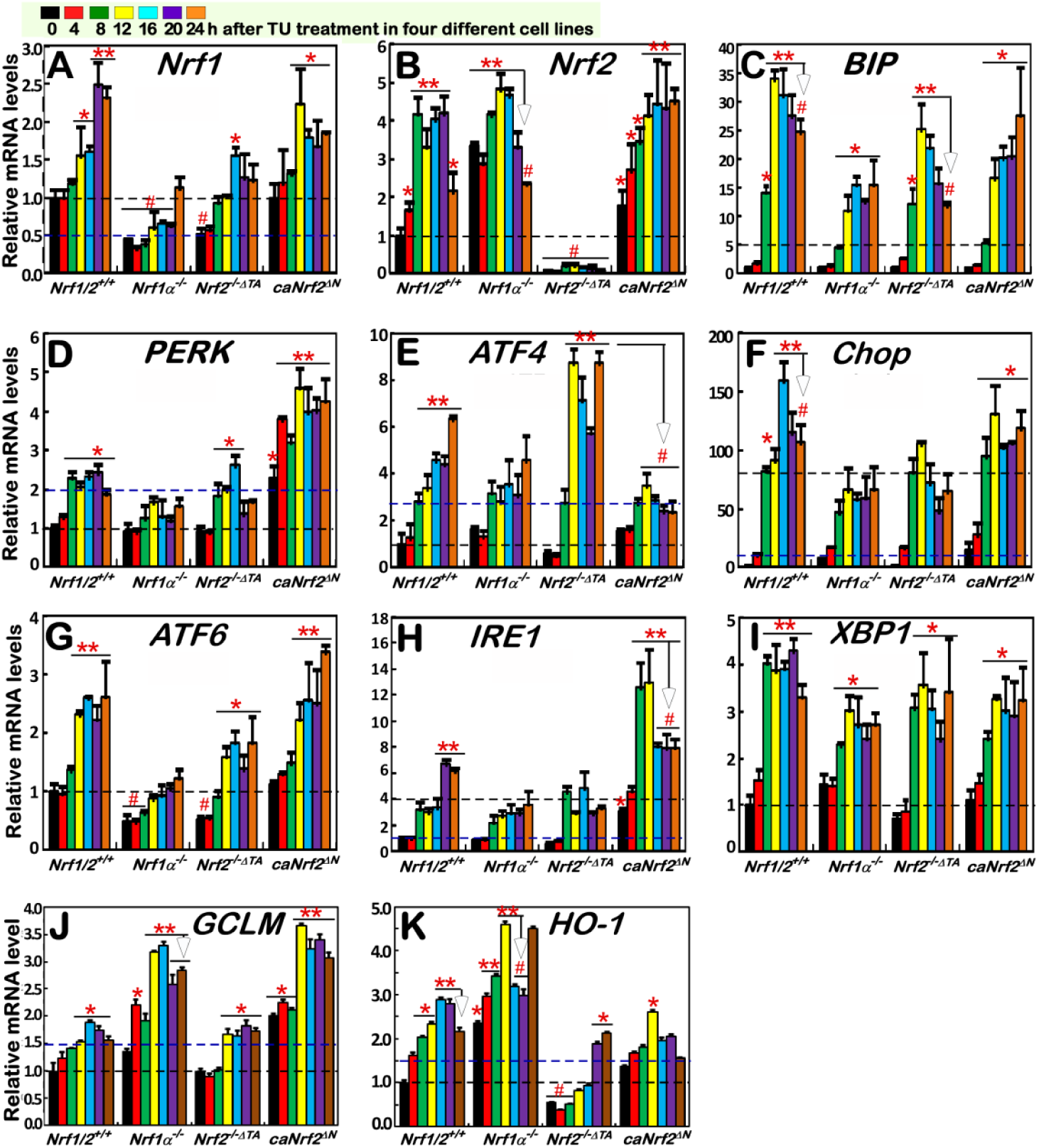
TU-inducible changes in the mRNA expression of distinct responsive genes. Distinct cell lines of *Nrf1/2*^*+/+*^, *Nrf1α*^*−/−*^, *Nrf2* ^*−/−δTA*^ and *caNrf2*^*δN*^ were treated with 2 μg/ml TU for the indicated times from 0 to 24 h. Then, TU-inducible mRNA expression of distinct responsive genes was determined by real-time qPCR. These genes include *Nrf1* (A), *Nrf2* (B), *BIP/GRP78* (C), *PERK* (D), *ATF4* (E), *Chop* (F), *ATF6* (G), *IRE1* (H), *XBP1* (I), *GCLM* (J) and *HO-1* (K). Significant statistical decreases were indicated with #, *p*<0.01, whereas significant increases were represented by \**p*<0.01 and \*\**p*<0.001.

Further qRT-PCR analysis of *Nrf1/2*^*+/+*^ cells revealed that a TU-stimulated increase in the mRNA expression of *BIP* (as a classic marker of ER stress-induced UPR) from 8 h to 24 h following treatment, which occurred with a peak of TU induction at 12 h (Figure 3C). By contrast, such TU-inducible mRNA expression of *BIP* was substantially suppressed and postponed in *Nrf1α*^*−/−*^ cells (with hyper-expression of Nrf2). Also, *Nrf2*^*−/−ΔTA*^ cells only gave a modest reduction of TU-inducible *BIP* expression, with a lowered peak at 12 h and a subsequent downward course to 24 h after treatment. Collectively, these indicate that Nrf1 is required for transcriptional regulation of UPR-target *BIP* gene, while Nrf2 is partially involved in this response to TU. However, *caNrf2*^*ΔN*^ (acting as a constitutive CNC-bZIP activator) also caused a partial decrease of TU-inducible *BIP* expression (Figure 3C). This decrease is attributable to a deletion of the Keap1-binding Neh2 domain from Nrf2 (to yield *caNrf2*^*ΔN*^), albeit the nuclear function of Neh2 is not yet identified to date.

Analysis of the PERK-eIF2α-ATF4-Chop response pathway unraveled that TU treatment of *Nrf1/2*^*+/+*^ cells caused distinct time-dependent increases in mRNA levels of *PERK, ATF4* and *Chop* from 8 h to 24 h (Figure 3D,3E,3F), which occurred with their respective peaks at 8 h, 24 h and 16 h. Of note, TU-inducible expression of *PERK* was substantially prevented in *Nrf1α*^*−/−*^ cells (with high expression of Nrf2), but appeared to be almost unaffected by *Nrf2*^*−/−ΔTA*^, when compared with those obtained from *Nrf1/2*^*+/+*^ cells (Figure 3D). This suggests that Nrf1α, but not Nrf2, is essential for transcriptional regulation of *PERK* in response to TU. Nevertheless, *caNrf2*^*ΔN*^ gave rise to rather significant increments in basal and TU-induced mRNA expression of *PERK* (Figure 3D). Hence, this implies that, once Nrf2 is localized in the nucleus, its Neh2 domain might serve as a putative dominant *trans*-repressor of the *PERK* gene. Known as a direct substrate of PERK, eIF2α is phosphorylated in the ER response to TU [4] and then contributes to selective translation of ATF4 (Figure 1). Herein, qRT-PCR showed that a time-dependent increment in *ATF4* mRNA levels was induced in *Nrf1/2*^*+/+*^ cells that had been treated for 8 h to 24 h with TU, which was peaked at 24 h (Figure 3E). Such late stages of TU-inducible *ATF4* response after 16-h treatment were blocked in *Nrf1α*^*−/−*^ or *caNrf2*^*ΔN*^ cells. Conversely, *Nrf2*^*−/−ΔTA*^ led to a remarkable accelerated promotion of *ATF4* expression induced by TU, with an early peak at 12 h (Figure 3E). These results indicate that Nrf2, but not Nrf1α, acts as a dominant trans-repressor of *ATF4* gene in cellular response to TU. Further qRT-PCR analysis of *Chop* (as a downstream target gene of ATF4) revealed that its TU-inducible mRNA expression was markedly blocked by *Nrf1α*^*−/−*^, and also partially inhibited by *Nrf2*^*−/−ΔTA*^, but seemed to be unaffected by *caNrf2*^*ΔN*^ (Figure 3F). Thereby, it is inferable that Nrf1α contributes primarily to transcriptional regulation of *Chop* gene, albeit Nrf2 is partially involved in this process.

Close examinations of other two ER signaling arms ATF6 and IRE1 showed distinct time-dependent induction of their mRNA expression by TU treatment of *Nrf1/2*^*+/+*^ cells, respectively with different peaks at 12 h or 20 h (Figure 3G, 3H). Such significantly TU-induced increases in *ATF6* mRNA expression were substantially suppressed in *Nrf1α*^*−/−*^ cells, and also partially inhibited by *Nrf2*^*−/−ΔTA*^, but exhibited almost no changes in *caNrf2*^*ΔN*^ cells (Figure 3G). This indicates that Nrf1α is required for regulating transcriptional expression of *ATF6* gene induced by TU, while Nrf2 only makes a minor contribution to this response. By contrast with *Nrf1/2*^*+/+*^ cells, induction of *IRE1* mRNA expression by TU was obviously prevented in *Nrf1α*^*−/−*^ cells (with hyper-expression of Nrf2), but not recovered in *Nrf2*^*−/−ΔTA*^ cells (Figure 3H), implying a major contribution of Nrf1α, rather than Nrf2, to *IRE1* transcriptional expression. As such, the late-stage induction of *IRE1* after 20 h TU treatment was also reduced by *Nrf2*^*−/−ΔTA*^. Contrarily, *caNrf2*^*ΔN*^ led to rather significant elevations in basal and TU-inducible *IRE1* mRNA levels, with an early higher peak that stimulated at 8-h TU treatment and maintained to 12 h, after being abruptly lowered to similar levels to the late-stage induction of *Nrf1/2*^*+/+*^ cells (Figure 3H). These suggest that dual opposing roles of Nrf2 in the *IRE1* transcriptional response to TU may depend on the presence of distinct functional domains within this CNC-bZIP factor and/or their biochemical modifications by the putative signaling, albeit the detailed mechanisms are unknown. Besides, transcriptional expression of *XBP1* mRNA as a direct substrate of IRE1 was further analyzed by qRT-PCR. The results disclosed that an accelerated increase in *XBP1* mRNA expression to a 4-fold maximal value was induced by TU after 8-h treatment of *Nrf1/2*^*+/+*^ cells (Figure 3I). By comparison with wild-type, TU-inducible *XBP1* expression was significantly decreased in *Nrf1α*^*−/−*^ cells, and also partially reduced by *Nrf2*^*−/−ΔTA*^ or *caNrf2*^*ΔN*^. Hence, it is postulated that Nrf1α and Nrf2 are involved in co-regulating the responsive expression of the *XBP1* gene to TU.

Next, examinations of ARE-driven genes by qRT-PCR revealed that TU treatment caused distinct time-dependent induction of *GCLM* and *HO-1* expression in *Nrf1/2*^*+/+*^ cells (Figure 3J,3K). By contrast, induction of *GCLM* and *HO-1* by TU was further significantly incremented in *Nrf1α*^*−/−*^ cells (retaining hyper-expression of Nrf2). Furtherly, *caNrf2*^*ΔN*^ also gave rise to substantial increases in basal and TU-induced mRNA levels of *GCLM*, rather than *HO-1* (Figure 3J,3K). Conversely, *Nrf2*^*−/−ΔTA*^ led to obvious decreases in basal and TU-induced mRNA levels of *HO-1*, but not *GCLM* (Figure 3J,3K). These imply that transcriptional responses of *HO-1* and *GCLM* to TU are mediated dominantly by Nrf2, albeit Nrf1 partially contributes to this antioxidant response. In addition, TU-inducible expression of *HO-1* is also partially reduced by *caNrf2*^*ΔN*^ (Figure 3K), implicating a possible attribution to the loss of the Neh2 domain from Nrf2. Taken altogether, our results demonstrate that transcriptional expression of *Nrf1* and *Nrf2*, as well as co-target genes *GCLM* and *HO-1*, is differentially induced by ER stressor, as accompanied by distinct activation of UPR signaling networks.

### Distinct contributions of Nrf1α and Nrf2 to protein expression of antioxidant genes induced by ER stressor

Here, we further determined changes in the protein expression of Nrf1 and Nrf2, along with downstream antioxidant genes *GCLM* and *HO-1*, in different genotypic cellular responses to TU stress stimulation for distinct lengths of time from 4 h to 24 h. Western blotting of *Nrf1/2*^*+/+*^ cells that had been treated TU (as a specific inhibitor of oligosaccharyl transferases to block N-glycosylation of newly-synthesized polypeptides) revealed that abundance of the full-length Nrf1 glycoprotein-A was gradually decreased from 8-h to its disappearance (Figure 4A1). Instead, the abundances of deglycosylated and processed Nrf1 protein-C/D were relatively incremented as the TU treatment time was increased. By sharp contrast, all these Nrf1-derived isoforms were completely abolished by specific knockout of *Nrf1α*^*−/−*^ (Figure 4B1). In addition, Nrf1α-derived protein levels were also modestly influenced by *Nrf2*^*−/−ΔTA*^ or *caNrf2*^*ΔN*^, because both mutants gave rise to an obviously-accelerated disappearance of Nrf1 glycoprotein within 4 h to 8 h of TU treatment (Figure 4C1,4D1), when compared with its presence in *Nrf1/2*^*+/+*^ cells (Figure 4A1). Conversely, this disappearance of Nrf1 glycoprotein was, rather, replaced by accelerated abundances of Nrf1 protein-C/D in *caNrf2*^*ΔN*^ cells (Figure 4D1). Together, these results demonstrate that glycosylation, deglycosylation and proteolytic processing of Nrf1 (associated with the ER) are regulated by TU-induced stress response signaling, part of which may be mediated by Nrf2.

**Figure 4.**
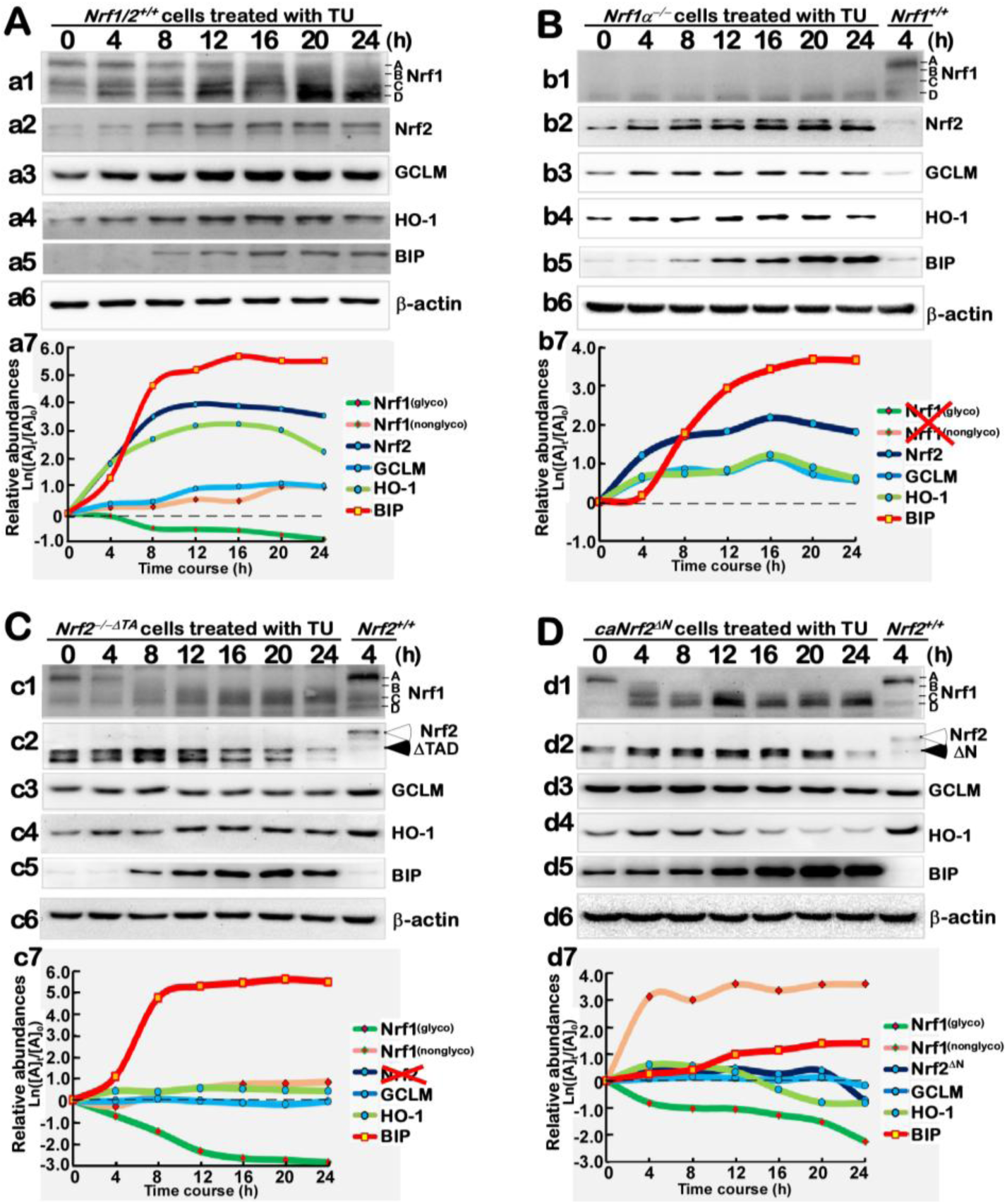
TU-inducible changes in the protein expression of antioxidant genes. Distinct cell lines of *Nrf1/2*^*+/+*^ (A), *Nrf1α*^*−/−*^ (B), *Nrf2* ^*−/−δTA*^ (C) and *caNrf2*^*δN*^ (D) were treated with 2 μg/ml TU for the indicated times from 0 to 24 h. The TU-inducible changes in the protein expression of distinct responsive genes were determined by Western blotting with the indicated antibodies against Nrf1, Nrf2, GCLM, HO-1, BIP/GRP78 or β-actin. The intensity of blots representing different protein expression levels were also quantified by the Quantity One 4.5.2 software and shown graphically.

Indeed, it is true that a gradual increment in Nrf2 protein abundances resulted from 4-h to 24-h TU-treatment of *Nrf1/2*^*+/+*^ cells (Figure 4A2). More intriguingly, a major processed isoform of Nrf2 was gradually incremented with the increasing time of TU treatment of *Nrf1α*^*−/−*^ cells (Figure 4B2). Similar observations was also represented in *caNrf2*^*ΔN*^, but not *Nrf2*^*−/−ΔTA*^ cells (*c.f.* Figure 4D2 with 4C2). Hence, it is inferable that a putative proteolytic processing of Nrf2 may occur through within its N-terminal Neh2 domain, albeit the detailed mechanism requires to be elucidated.

Further examinations of antioxidant protein expression unraveled that distinct time-dependent increments of GCLM and HO-1 were significantly induced by TU treatment in *Nrf1/2*^*+/+*^ cells (Figure 4A3,4A4). Similar induction of GCLM and HO-1 by TU was also observed in *Nrf1α*^*−/−*^ cells (Figure 4B3,4B4), but was obviously reduced by *Nrf2*^*−/−ΔTA*^ (Figure 4C3, 4C4), with compared with the controls from *Nrf1/2*^*+/+*^ cells. However, *caNrf2*^*ΔN*^ cells displayed almost no changes in GCLM protein (Figure 4D3); this was accompanied by a modest decrease in HO-1 expression (Figure 4D4). Collectively, these results demonstrate that, although Nrf2 is negatively regulated by Nrf1, the former Nrf2 makes a major contribution to regulating expression of *GCLM* and *HO-1* genes, possibly through its N-terminal Neh2 domain.

Next, Western blotting of the ER stress-responsive chaperone BIP showed its protein abundance was increased in a time-dependent manner from 4 h to 24 h of TU treatment of *Nrf1/2*^*+/+*^ cells (Figure 4A5). Interestingly, remarkable increments in BIP protein levels were presented in TU-treated *Nrf1α*^*−/−*^ and *Nrf2*^*−/−ΔTA*^ cells (Figure 4B5,4C5), albeit its mRNA expression levels were lowered to different extents in these two cell lines (Figure 3C). In addition, basal and TU-stimulated BIP abundances were also strikingly incremented in *caNrf2*^*ΔN*^ cells (Figure 4D5). Overall, these indicate that Nrf1α and Nrf2 are not essential for mediating BIP protein expression, albeit both CNC-bZIP factors are involved in this chaperone transcriptional response to TU, besides antioxidant response to this ER stressor.

### Distinct involvements of Nrf1α and Nrf2 in differential expression of TU-induced ER stress responsive genes

Clearly, it is known that the chaperone BIP protein is a key sensor to ER stress induced by TU, so that expression of its cognate gene as a direct effector is activated in UPR [1-4]. Such an ER stress model was also successfully constructed as described above. Herein, we further examined whether Nrf1α and Nrf2 are required for crucial protein expression for the UPR signaling cascades. As anticipated, the phosphorylated protein levels of PERK were significantly increased following 12 h to 16 h of TU treatment of *Nrf1/2*^*+/+*^ cells (Figure 5A2). The induction of p-PERK by TU appeared to be prevented in *Nrf1α*^*−/−*^ and *Nrf2*^*−/−ΔTA*^ cells (Figure 5B2, 5C2). By contrast, *caNrf2*^*ΔN*^ led to an early modest induction of p-PERK by TU at 4 h to 8 h following treatment (Figure 5D2). These suggest that Nrf1α and Nrf2 may be required for regulation of the PERK signaling response to TU.

**Figure 5.**
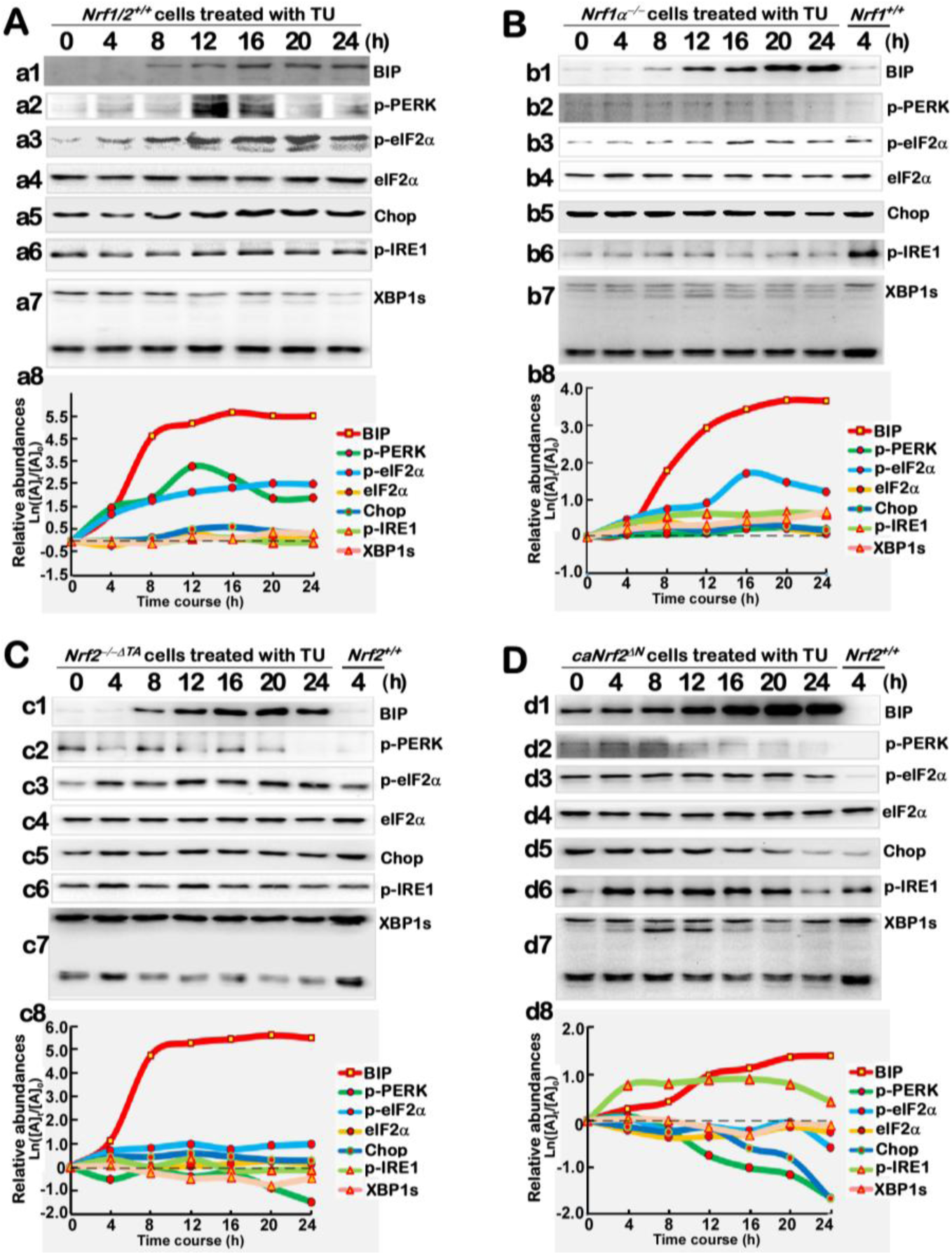
TU-inducible changes in the protein expression of ER stress-responsive genes. Distinct cell lines of *Nrf1/2*^*+/+*^ (A), *Nrf1α*^*−/−*^ (B), *Nrf2* ^*−/−δTA*^ (C) and *caNrf2*^*δN*^ (D) were treated with 2 μg/ml of TU for the indicated times from 0 to 24 h. The TU-inducible changes in the protein expression of distinct responsive genes were determined by Western blotting with the indicated antibodies against BIP/GRP78 (as a positive reference, that was duplicated from Figure 4), p-PERK, p-eIF2α, eIF2α, Chop, p-IRE1 or XBP1. The intensity of blots representing different protein expression levels were also quantified by the Quantity One 4.5.2 software and shown graphically.

Meanwhile, the total protein abundances of eIF2α (as a main substrate of p-PERK to yield p-eIF2α) were almost unaltered by TU in all the above-described four cell lines (Figure 5A4, 5B4, 5C4, 5D4). In addition to the basal eIF2α auto-phosphorylation, its TU-inducible phosphorylation was also enhanced as the time of treatment was increased from 4 h to 16 h, and then maintained until 24 h after treatment of *Nrf1/2*^*+/+*^ cells (Figure 5A3). However, TU-induced eIF2α phosphorylation was almost unaffected in *Nrf1α*^*−/−*^ cells (Figure 5B3), when compared with the control of 4-h TU-treated *Nrf1/2*^*+/+*^ cells. Conversely, induction of eIF2α phosphorylation by TU was partially recovered in *Nrf2*^*−/−ΔTA*^ cells (Figure 5C3). These results indicate that Nrf1α and Nrf2 may contribute to positive and negative regulation of eIF2α induction by TU, respectively. Yet, *caNrf2*^*ΔN*^ gave rise to an increase in basal eIF2α auto-phosphorylation and its TU-inducible phosphorylation by 20 h of treatment (Figure 5D3), by comparison with the control of 4-h TU-treated *Nrf1/2*^*+/+*^ cells. This intriguing data implicates that eIF2α may be negatively regulated by the N-terminal Neh2 domain of Nrf2, besides its transactivation domains (because both lacked in *caNrf2*^*ΔN*^ and *Nrf2*^*−/−ΔTA*^ cells, respectively).

Western blotting examination of Chop, as an effector of the PERK-eIF2α-ATF4 signaling pathway, revealed that a modest increment in Chop protein levels resulted from 8 h to 24 h of TU treatment of *Nrf1/2*^*+/+*^ cells (Figure 5A5). By contrast, almost no changes in Chop abundances were determined in *Nrf1α*^*−/−*^ and *Nrf2*^*−/−ΔTA*^ cells, albeit both lines had been treated with TU (Figure 5B5,5C5). However, basal Chop abundance was obviously elevated by *caNrf2*^*ΔN*^ and appeared to be almost unaffected by TU within 12 h after this treatment, but thereafter decreased gradually to a similar level to the control of 4-h TU-treated *Nrf1/2*^*+/+*^ cells (Figure 5D5). Together, these results indicate that Chop is co-regulated by Nrf1α and Nrf2, but the latter Nrf2 may also contribute to negative regulation of Chop possibly by its Neh2 domain, in the ER-to-nuclear response to the long-term TU-induced stress.

Next, distinct time-dependent expression of the IRE1/ATF6-XBP1 signaling molecules was examined by Western blotting. The results unraveled that, since the existing auto-phosphorylation of IRE1 has been at a considerable high level, TU only induced a modest increase in its phosphorylated abundance following 12-h treatment of *Nrf1/2*^*+/+*^ cells (Figure 5A6). Such low induction of IRE1 by TU appeared to be almost abolished in *Nrf1α*^*−/−*^ cells (Figure 5B6, when compared to the control of 4-h TU-treated *Nrf1/2*^*+/+*^ cells), but also was only slightly recovered from 4 h to 12 h of TU treatment of *Nrf2*^*−/−ΔTA*^ cells (Figure 5C6). However, *caNrf2*^*ΔN*^ cells raised a rather higher induction of phosphorylated IRE1 by TU (Figure 5D6). Together, these indicate that Nrf1α and Nrf2 are involved in regulating the IRE1 signaling response to TU. In addition, it is important to note that the intact XBP1u mRNA, though as a direct substrate of IRE1 to yield a specifically-spliced XBP1s (besides its mRNA decay), is also transcriptionally regulated by ATF6 signaling in ER stress response [[1-4], Figure 1]. Therefore, Western blotting showed that a few of processed XBP1s bands were observed in *Nrf1/2*^*+/+*^ cells (Figure 5A7), and also obviously enhanced by TU in *Nrf1α*^*−/−*^ or *caNrf2*^*ΔN*^ cells (Figure 5B7, 5D7), but not determined in *Nrf2*^*−/−ΔTA*^ cells (Figure 5C). Conversely, the putative XBPu protein was rather reduced by *Nrf2*^*−/−ΔTA*^ only. These results implicate that Nrf2 may contribute to regulating XBP response to TU, albeit the detailed mechanism needs to be further elucidated.

### No induction of proteasomal (*PSM*) subunit genes regulated by Nrf1 in the response to TU stressor

As described by us and others [22, 24, 29], it is rather clear that Nrf1, but not Nrf2, exerts an important biological role in the transcriptional expression of all *PSM* genes. Such transcriptional regulation of *PSM* genes by Nrf1 is also accompanied by induction of all three classic ER-driven stress response signaling pathways mediated by PERK1, IRE1 and ATF6, along with the proteasomal compensatory response to limited extents of proteasome inhibitors [24]. Here, we further examined whether TU-stimulated Nrf1 and Nrf2 are required for the ER-to-nuclear signaling responses to transactivate *PSM* genes. The results revealed that none of all six *PSM* mRNA expression levels was induced by TU in *Nrf1/2*^*+/+*^ cells (Figure 6A). Contrarily, *PSMA1* and *PSMB7* mRNA levels were obviously down-expressed in TU-treated *Nrf1/2*^*+/+*^ cells (Figure 6A1, 6A5).

**Figure 6.**
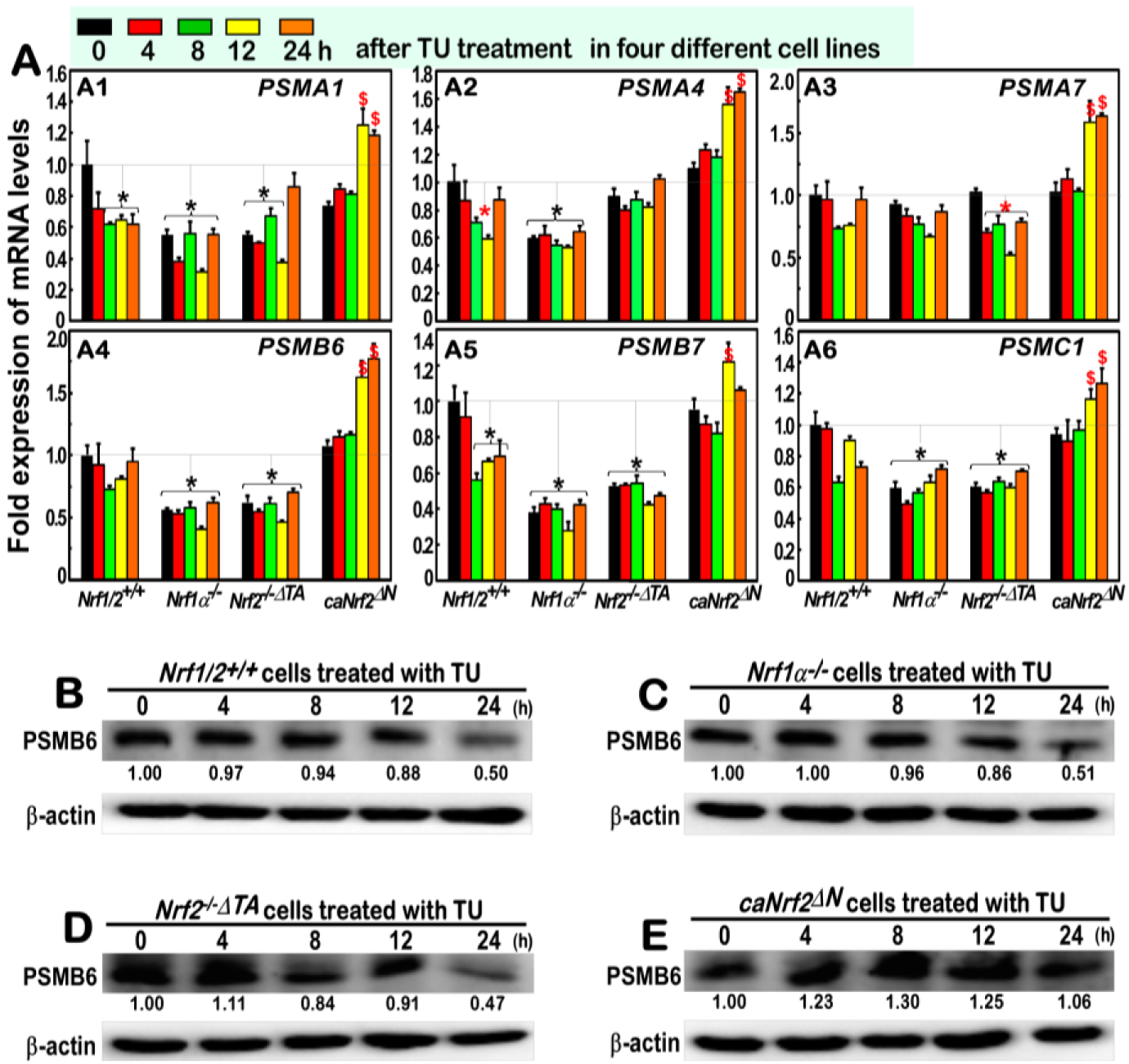
No obvious induction of some examined proteasomal genes by TU. Distinct cell lines of *Nrf1/2*^*+/+*^, *Nrf1α*^*−/−*^, *Nrf2* ^*−/−δTA*^ and *caNrf2*^*δN*^ were treated with 2 μg/ml TU for the indicated times from 0 to 24 h. The TU-inducible mRNA levels of some proteasomal genes, including *PSMA1, PSMA4, PSMA7, PSMB6, PSMB7 and PSMC1*, were determined by real-time qPCR (A), and Western blotting with indicated antibodies against PSMB6 or β-actin in different cell lines *Nrf1/2*^*+/+*^ (B), *Nrf1α*^*−/−*^ (C), *Nrf2* ^*−/−δTA*^ (D) and *caNrf2*^*δN*^ (E). The intensity of blots representing PSMB6 was also quantified by the Quantity One 4.5.2 software, and shown on the *bottom*.

Further qRT-PCR analysis of *Nrf1α*^*−/−*^ cells unraveled that the basal mRNA expression levels of *PSMA1, PSMA4, PSMB6, PSMB7* and *PSMC1*, but not *PSMA7*, were significantly down-regulated and also unaffected by TU stimulation (Figure 6A). Similar down-regulation of *PSMA1, PSMB6, PSMB7, PSMC1*, but not *PSMA4* or *PSMA7*, was detected in *Nrf2*^*−/−ΔTA*^ cells. However, all these six *PSM* genes were only induced by the long-term exposures of TU in *caNrf2*^*ΔN*^ cells from 12 h to 24 h. Moreover, Western blotting of *PSMB6* disclosed that its protein abundance was decreased by TU, with the increasing treatment time of *Nrf1/2*^*+/+*^ cells (Figure 6B). Similarly TU-triggered decreases in *PSMB6* were almost unaltered by *Nrf1α*^*−/−*^ or *Nrf2*^*−/−ΔTA*^ cells (Figure 6C,6D), but markedly reversed by *caNrf2*^*ΔN*^ cells with a modest increase (Figure 6E). Collectively, distinct contributions of Nrf1α and Nrf2 to basal, but not TU-inducible, expression of some *PSM* genes are demonstrated. Yet, Nrf2 might exert opposing roles in this event, depending on its functional domains within different responsive contexts.

## Discussion

In the present study, it is demonstrated that there exists a bi-directional cross-talk between UPR-triggered signaling and ARE-driven cytoprotective responses to the ER stressor TU (Figure 7). Importantly, we have also demonstrated the evidence that opposite roles of Nrf1 and Nrf2 are unified to coordinate different cellular responses to TU, leading to differential activation of ER-driven stress signaling networks. Of note, loss of Nrf1 down-regulates expression of antioxidant, detoxification and particularly proteasomal genes, leading to severe oxidative stress and concurrently ER stress [27, 34]. The latter pathophysiological event is primarily attributable to disruption of protein folding within the ER and dysfunction of ERAD, such that unfolded and misfolded proteins, along with oxidized and damaged proteins, are accumulated within the ER. Consequently, the canonical UPR signaling pathways are activated by PERK, IRE1 and ATF6 (Figures 1 and 7), in order for the ER adaptive remodeling to diminish loading of nascent polypeptides, remove aberrant folded proteins, and then restore itself biological function of this organelle [35, 36]. This notion is supported by the evidence showing that endogenous ER stress signaling to activate UPR occurred in the steatotic hepatocytes with a homozygous knockout of *Nrf1*^*−/−*^ [27], but not *Nrf2*^*−/−*^ [12] and similar ER stress-inducible response was further enhanced by proteasomal inhibition of the heterozygous *Nrf1*^*+/−*^ livers, when compared with wild-type [27]. Thereby, Nrf1 plays an essential role in maintaining the ER homeostasis, but its functional loss results in the ER transformation and proliferation of *Nrf1*^*−/−*^ cells in conditional knockout mice, that are spontaneously developed with non-alcoholic steatohepatitis (NASH) and liver cancer [34]. Such phenotypes should be embodied as a pathological consequence of chronic stimulation of ER stress and prolonged activation of UPR signaling, as occurred concomitantly with severe oxidative stress, which together lead to carcinogenesis ultimately [37-39].

**Figure 7.**
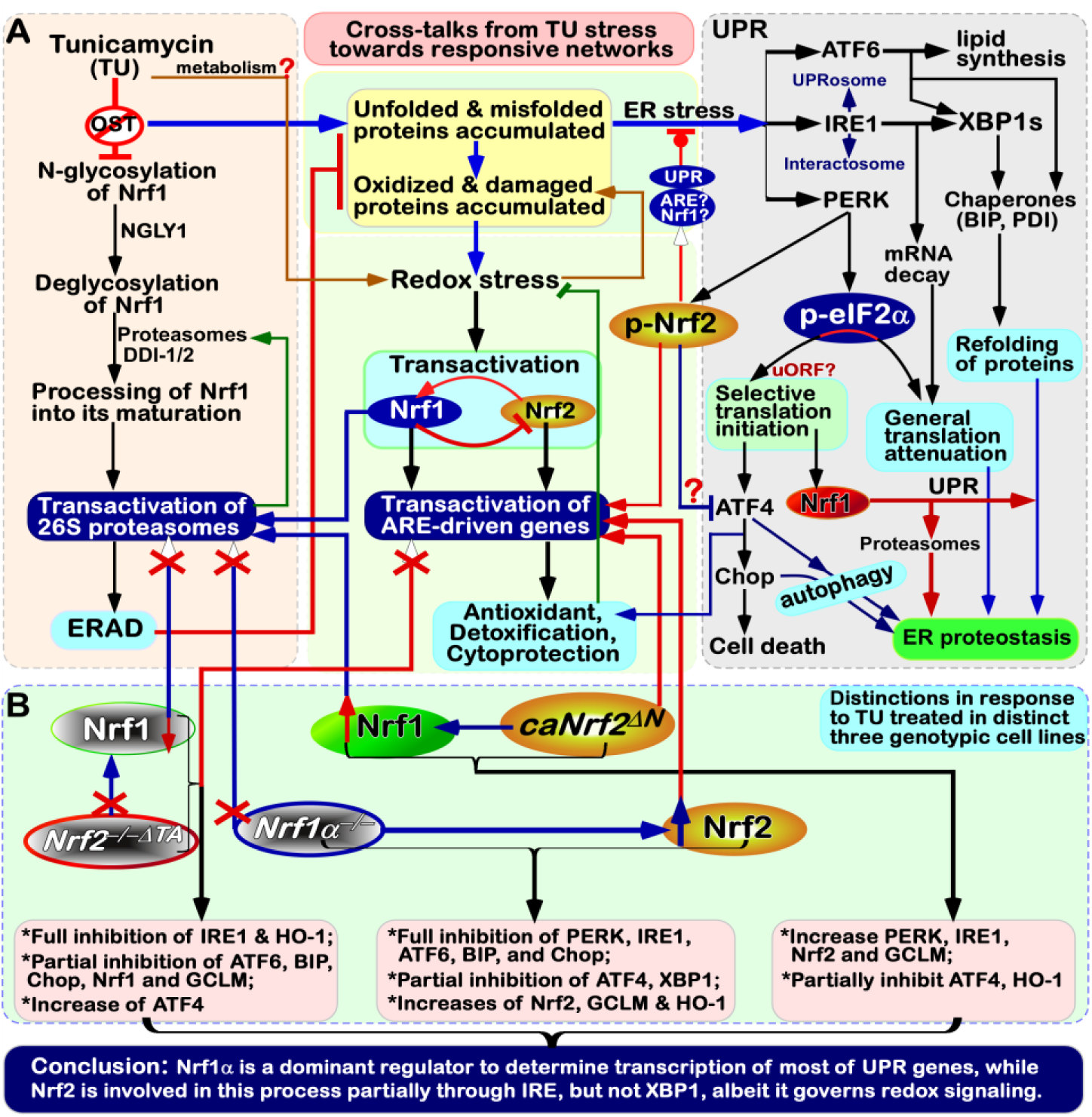
Cross-talks from TU stress signaling to different ER-to-nuclear responsive networks. (A) There exist multiple cross-talks from TU stress signaling to different ER-to-nuclear responsive networks. Inhibition of Nrf1 N-glycosylation by TU leads to subsequent blockage of its deglycosylation and proteolytic processing to yield a mature CNC-bZIP factor, before transactivating proteasomal genes. Such disruption of Nrf1-mediated proteasomal degradation (e.g., ERAD) leads to aberrant accumulation of oxidized, damaged and misfolded proteins, in addition to TU-led non-glycosylated and unfolded proteins. These together result in classic ER stress along with redox stress. Of note, distinct and even opposing roles of Nrf1 and Nrf2, as well as both inter-regulatory effects on cytoprotection against ER redox stress, are integrally unified in different cellular signaling responses within distinct gene-regulatory networks. (B) Our evidence demonstrates that Nrf1 acts as a dominant regulator of most of UPR-target genes, apart from its negative regulation of Nrf2. By contrast, Nrf2, besides as a direct substrate of PERK, is also involved in this response to TU partially through the IRE1 signaling pathway, but not by its downstream XBP1. Although Nrf2 governs transcription of Nrf1 and their co-target antioxidant genes, it also contributes to negative regulation of ATF4, that is selectively translated by the eIF2α-based machinery. As described for ATF4, Nrf1 is predicted to contain an upstream open reading frame (uORF) within its full-length mRNA transcript.

Notably, Nrf1 is a moving transmembrane protein with dynamic membrane-topologies (that are distinctive from those of PERK, IRE1 and ATF6 within and around the ER, as deciphered in Figure 1), and its topovectorial processes determine its post-synthetic modification and its *trans*-activity to mediate target *PSM* genes [24, 32]. Interestingly, our evidence has also been provided showing that the canonical UPR signaling by PERK, IRE1 and ATF6 to differential expression of distinct responsive genes is activated by classic ER stressor TU; this is accompanied by transcriptional expression of Nrf1 and Nrf2, as well as their co-target *GCLM* and *HO-1* (Figure 7). This appears to raise a paradoxical question about Nrf1, because N-glycosylation of this CNC-bZIP protein (that is newly synthesized in the ER lumen) is sufficiently blocked by TU, with secondary inhibition of its ensuing deglycosylation and proteolytic processing to yield an N-terminally-truncated mature factor in close proximity to membranes. This inhibitory effect of TU on Nrf1 is also further supported by no induction of all examined *PSM* genes regulated by Nrf1, rather than Nrf2, leading to ERAD dysfunction so as to exacerbate accumulation of those misfolded, oxidized and damaged proteins by this stressor TU. In a feedback regulatory response to mitigate this deteriorating ER stress, Nrf1 is transcriptionally activated by Nrf2. This is based on the fact that Nrf1 is identified as a direct target gene of Nrf2 [31] and also Nrf2, as a direct substrate of PERK, is activated by TU [10-12]. Further translational expression of Nrf1 could also be selectively initiated by a putative mechanism accounting for ATF4, c-Myc and C/EBP [40-42] (Figure 7), which can enable their translational expression to switch from their upstream open reading frames (uORF) to their main open reading frames (mORF), in the cellular response to ER stress [8]. Such being the case, it is hence postulated that such uORF (and/or its products) could act as a molecular pathophysiological switch driving carcinogenesis or other degenerative diseases.

Although the above relevant detailed mechanisms warrant to be further elucidated, our evidence has been here presented demonstrating that Nrf1 is significantly increased at both its transcriptional and translational expression levels in TU-stimulated of *Nrf1/2*^*+/+*^ cells, particularly of *caNrf2*^*ΔN*^ cells (with an enhanced constitutive expression of Nrf2). This implies that a portion of non-glycosylated Nrf1 and its subsequent processed isoforms are incremented in a feedback regulatory circuit. However, it is inferable that a fraction of such non-glycosylated and processed proteins of Nrf1 (that are generated during the TU-stress conditions) only has a weak activity to mediate target genes, relative to another distinctive fraction of its deglycosylated and processed proteins (that are generated in TU-untreated cells). In fact, this point is convincingly evidenced by inhibition of peptide:N-glycosidase (PNG-1/NGLY1, an evolutionarily conserved deglycosylation enzyme) to inactivate Nrf1 and thus down-regulate expression of its target *PSM* genes, so as to potentiate cytotoxicity of proteasomal Inhibitors [43]. Besides, the NGLY1-NRF1 pathway is revealed to exert an additional novel functions in mitochondrial homeostasis and inflammation pathogenesis [44]. Similar inactivation of Nrf1 to disrupt deglycosylation of misfolded N-glycosylated proteins has been recently unveiled by loss-of-function of NGLY1, a homozygous mutant condition that has been characterized by a complex neurological syndrome, hypo- or alacrimia, and elevated liver transaminases [45, 46]. Similarly, the functional analysis of a *Drosophila* model of NGLY1 deficiency to its CNC inactivation provides insight into therapeutic approaches [47]. Moreover, additional homologue SKN-1A associated with the ER of *Caenorhabditis elegans* also mediates a cytoplasmic unfolded protein response and promotes longevity [48]. This biological event occurs to be regulated by the amino acid sequence editing of SKN-1A by NGLY1 from its glycosylated Asn (-N-S/T-) into acidic Asp (-D-S/T-) residues within this protein, in order to control expression of *PSM* genes against proteotoxic stress [49]. Overall, these supportive findings are prime to confirm our prior work on Nrf1 glycosylation and deglycosylation to finely tune its transactivation activity [20, 33, 50, 51].

Moreover, it should also be noted that the dynamic membrane-topology of Nrf1 to determine its post-synthetic processing and its transactivation activity might be modulated by cholesterol within membranes in the ER-to-nuclear signaling response to cellular stress. This viewpoint is attributable to direct binding of Nrf1 to cholesterol by its five cholesterol-recognized amino acid consensus (CRAC) sites within this CNC-bZIP protein [32, 51, 52]. Recently, the ER-associated Nrf1 is also identified as a vital sensor to cholesterol within membranes [53]. These collective findings demonstrate that Nrf1 plays a central role in maintaining cholesterol homeostasis by a negative feedback regulatory loop against an additional ER membrane-bound SREBP2 (sterol response element binding protein). In reality, the UPR signaling mediated by SREBP (and ATF6) has also been showed a linkage to a lipid stress response [54-56]. Activation of SREBP signaling occurs only after being translocated into the Golgi apparatus, in which it is subject to sequential cleavages by Site-1 and Site-2 proteases in a similar fashion to the case of ATF6 (Figure 1). However, Nrf1 is neither translocated into the Golgi apparatus and nor cleaved by both Site-1 and Site-2 proteases [33, 50]. As a master of fact, the *bona fide* activation of Nrf1 occurs only after its dynamic flipping out of ER membranes and then by its topology-regulated juxtamembrane proteolytic processing by cytosolic proteasomes and DDI-1/2 proteases to yield a mature CNC-bZIP factor [8, 24, 32, 57]. Besides, SREBP1 was also identified as a direct upstream regulator of Nrf1, which is involved in the mTORC1 signaling response to insulin and epidermal growth factor [23]. Yet, it remains unknown whether there exists a coordinated cross-talk between oxidative stress response and lipid-coupled UPR. As such, it is plausible that Nrf1 senses cholesterol changes in the vicinity of ER membranes, and relevant signals are integrated and transduced to down-regulate cognate target genes. Meanwhile, the cytoprotective responsive signaling to ER stress enables Nrf1 to be selectively translated by the putative uORF existing within the first exon of its full-length transcripts and/or processed by its topology-regulated juxtamembrane proteolysis. Such a unique mechanism is definitely distinctive from that accounts for both ATF6 and SREBP1/2 factors in the ER stress responses to unfolded proteins and overloaded lipids.

Here, it is of a crucial significance to reveal that specific knockout of *Nrf1α*^*−/−*^ leads to aberrant hyper-expression of Nrf2 and cognate target *GCLM and HO-1*, as companied by down-expression of most UPR-target genes (referenced in this study). Conversely, *Nrf2*^*−/−ΔTA*^ down-regulates its target genes *GCLM, HO-1 and Nrf1*, with a concurrent inhibition of some UPR-target genes, such as *IRE1, ATF6, BIP* and *Chop*. However, *ATF4* is strikingly up-regulated by inactivation of Nrf2, while their upstream *PERK* expression is almost unaffected. These findings, together with our previous work [24, 31], demonstrate that Nrf1α does not only dictate specific functioning of Nrf2 to be exerted in antioxidant responses, and also act as a dominant player in the regulation of most UPR-target genes (that contain the ARE sequences within their promoter regions, as listed in Table S1). Also, Nrf2 is partially involved in mediating the expression of some UPR genes by controlling transcription of *Nrf1* and *IRE1*, but not *XBP1*. However, expression of *ATF4* is down-regulated by *caNrf2*^*ΔN*^, albeit with up-regulation of *PERK* (and *IRE1*). This implies a putative mechanism for Nrf2 (as a direct substrate of PERK) to compete against the PERK-eIF2α-*ATF4* signaling (Figure 7). Furthermore, a partial inhibition of *HO-1* by *caNrf2*^*ΔN*^ also indicates a constitutive loss of its N-terminal Neh2 function as an extra unidentified nuclear transactivation domain after Nrf2 enters the nucleus.

## Conclusion

The evidence has been herein presented demonstrating a unity of distinct and even opposing roles of Nrf1 and Nrf2 in discrete cellular responses to ER stress induced by TU, hence leading to differential activation of the ER-to-nuclear signaling networks. Notably, there exist multiple cross-talks between UPR- and ARE-driven signaling cascades, aiming to maintain cellular redox, protein and lipid homeostasis. Altogether, these responses can make cell fate decisions to be modulated under ER stress conditions [58], as discovered for its homologous Skn-1 in *Caenorhabditis elegans* [18]. Importantly, Nrf1α (and/or its derivates) is a dominant regulator to determine the transcriptional expression of most of UPR-target genes, whereas Nrf2 is also involved in this ER-to nuclear response partially through IRE1 but not its substrate XBP1, albeit it is a central player in governing redox signaling networks.

## Materials and Methods

### Cell lines and Reagents

The human hepatocellular carcinoma HepG2 cells (i.e. *Nrf1/2*^*+/+*^) were obtained originally from the American Type Culture Collection (ATCC, Manassas, VA, USA). Three derived cell lines with knockout of *Nrf1α*^*−/−*^ or *Nrf2* ^*−/−δTA*^ and a constitutive activation of Nrf2 (i.e. *caNrf2*^*δN*^) were established by Qiu *et al* [31]. They were cultured in a 37°C incubator with 5% carbon dioxide, and allowed for growth in Dulbecco’s modified Eagle’s medium (DMEM) with 25 mmol/L high glucose, 10% (v/v) fetal bovine serum (FBS), 100 units/ml penicillin-streptomycin. TU (with MW 816.89) was purchased from Sangon Biotech Co., Ltd. (Shanghai, China). All pairs of primers used for qRT-PCR analysis are listed in Table S2.

### Cell viability

All four cell lines *Nrf1/2*^*+/+*^, *Nrf1α*^*−/−*^, *Nrf2* ^*−/−δTA*^ and *caNrf2*^*δN*^ were cultured for 24 h in DMEM containing 25 mmol/L glucose and 10% FBS. After reaching 70% of their confluences, they were then allowed for growth in fresh media with different concentrations of TU (at 0, 0.5, 1, 2, 4, or 8 μg/ml). For their time-course, they were also treated with 2 μg/ml of TU for different lengths of time (i.e. 0, 4, 8, 12, 16, 20, or 24 h). The cell viability was evaluated by using an MTT-based cell proliferation and cytotoxicity assay kit (Beyotime, China).

### The constitutive expression of ER stress-related genes in selected cell lines

Equal amounts of *Nrf1/2*^*+/+*^, *Nrf1α*^*−/−*^, *Nrf2* ^*−/−δTA*^, and *caNrf2*^*δN*^ were cultured in 6-well plates before being harvested in a lysis buffer [59]. Total cell lysates were subjected to protein separation by SDS-PAGE gels containing 8% polyacrylamide, followed by Western blotting with antibodies against Nrf1 (made in our laboratory) and Nrf2 (from ABCAM, Cambridge, UK) or β-Actin (from Zhong Shan Jin Qiao Co., Beijing, China). The β-Actin served as an internal control to verify amounts of proteins that were loaded in each of wells. Meantime, a portion of differential expression genes were identified by transcriptome sequencing, and their relative basal expression levels were also calculated and presented as fold changes (mean ± SD) in the RPKM (*R*eads *P*er *K*ilobase per *M*illion mapped reads). According to the Log2-based RPKM values against those determined from *Nrf1/2*^*+/+*^, the heatmaps for experimented cell lines were generated by using the MEV4.9 program.

### The mRNA expression of examined responsive genes to TU

After reaching 70% confluence of *Nrf1/2*^*+/+*^, *Nrf1α*^*−/−*^, *Nrf2* ^*−/−δTA*^ and *caNrf2*^*δN*^ cell lines grown in DMEM containing 25 mmol/L glucose and 10% FBS, they were treated for different time periods with 2 μg/ml of TU. Their total RNAs were extracted by using an RNA extraction kit (TIANGEN, China) and then subjected to reactions with reverse transcriptase (Promega, USA) to synthesize the single-strand cDNAs. Subsequently, relative mRNA expression levels of both ER stress-related and proteasomal genes in these cell lines were measured by RT-qPCR. This reaction was carried out in the GoTaq® real-time PCR detection systems, loaded on a CFX96 instrument (Bio-rad, USA). The results were analyzed by the Bio-Rad CFX Manager 3.0 software.

### The protein expression of examined responsive genes to TU

After reaching 70% confluence of *Nrf1/2*^*+/+*^, *Nrf1α*^*−/−*^, *Nrf2* ^*−/−δTA*^ and *caNrf2*^*δN*^ cell lines growth in DMEM containing 25 mmol/L glucose and 10% FBS, they were treated with 2 μg/ml of TU from distinct lengths of time. Their total lysates were separated by SDS-PAGE gels containing 10% polyacrylamide, followed by Western blotting with distinct primary antibodies against GCLM, HO-1, BIP, eIF2α, Chop, p-IRE1, XBP1 and PSMB6. These antibodies were purchased from Abcam, Inc (Cambridge, UK), except that p-PERK and p-eIF2 were from Cell Signaling Technology, Inc. (Massachusetts, USA). The β-Actin served as an internal control to verify amounts of proteins that were loaded in each of wells. The intensity of some immunoblots were quantified by the Quantity One 4.5.2 software and shown graphically.

### Statistical analysis

The ‘wet’ experimental data provided in this study were represented as the mean ± SD and were analyzed using the Student’s *t*-test or Fisher’s exact test, as appropriate. The resulting value of *p* < 0.05 was considered as a significant difference. In addition, statistical determination of the ‘dry’ sequencing analysis was described by Wang, *et al* [60].

### Data Availability

All data needed to evaluate the conclusions in the paper are present in this publication and/or the Supplementary Materials that can be found at xx. Additional data related to this paper may be requested from the authors.

## Author contributions

Y.-P.Z., S.H. and X.R. performed most of both experiments and bioinformatics, and collected relative data. Z.Z. and Z.F. also did some experiments. L.Q. made experimental cell lines used in this study. Y.-P.Z. prepared draft of this manuscript with most figures and supplementary tables. Y.Z. designed this study, analyzed all the data, helped to prepare all figures with two cartoons, wrote and revised the paper.

## Acknowledgments

The study was supported by the National Natural Science Foundation of China (NSFC, with key programs 91129703, 91429305 and project 81872336) awarded to Yiguo Zhang (Chongqing University, China).

