## Supplemental Table 1 and 2 for "A unity of opposites in between Nrf1- and Nrf2-mediated responses to the endoplasmic reticulum stressor tunicamycin"

### Supplementary tables:

**Table S1. ARE and AP1-binding sites in -5 kbp to TSS and also to TIS of ER-stress gene promoters**

| Gene ID | Name | ARE/EpRE (5'-TGAC/GnnnGC-3') | TRE/AP1 site (5'-TGAC/GTCA-3') |
|---|---|---|---|
| 3309 | BIP/GRP78 | TGGCGCAATCTCAGCTC (-4344 to -4328)<br>ATTTTGACCAGGCTGGT (-3811 to -3795)<br>TGGTGCGATCTCAGCTC (-2848 to -2832)<br>TAAGTGACTGTGCTTTG (-2480 to -2464)<br>GAGCTGAGATTGCACTA (-1339 to -1323) | CTCTTGAGTCACCAG (-2104 to -2090)<br>GTACTGAGTCACAGG (-2048 to -2034) |
| 9451 | PERK | GGTTTGAGTTCGCTCAT (-2728 to -2712)<br>TCTAGCAAACTCATATA (-1844 to -1828)<br>GCGTGCCAGGTCAGAGT (-719 to -703)<br>CCAATGAGAGAGCAAAC (+60 to +76) | |
| 2081 | IRE1 | ACCCGCCACCTCAGCCT (-4079 to -4063)<br>TGAGTGACTTGGCCGTG (-692 to -676)<br>AGTCTGACGCGGCAGGT (-370 to -354)<br>TGAGGCTCGGTCACCGC (+26 to +42) | |
| 22926 | ATF6 | TCTTGCTCTGTCACCCA (-3459 to -3443)<br>GAGCTGAGATGGCTCCA (-2054 to -2038)<br>GTTCTGAGATAGCCACG (-343 to -327) | |
| 1649 | CHOP | CACAGCTTGGTCATGTC (-4521 to -4505)<br>AAGGGCTACCTCAGTCA (-4384 to -4368)<br>AGGCGCCCTGTCACCCA (-2780 to -2764)<br>TCTCGCTCTGTCACCCA (-935 to -919)<br>AAGCTGAGTTGGCCAGG (+2219 to +2235) | CGCATGACTCACCCA (-242 to -228) |
| 7494 | XBP1 | TCCCTGACCGAGCTGGT (-4419 to -4403)<br>CACTGCAGCCTCAATCT (-4205 to -4189)<br>CTCAGCCTCCTCAGTAG (-3987 to -3971)<br>ATGTTGACCAGGCTGGT (-3901 to -3885)<br>CTGTTGACCAGGCTGGA (-2943 to -2927)<br>CTGGTGACAGAGCCTGA (-869 to -853)<br>AAATGCACGCTCATAGT (-701 to -685) | GGCATGAGTCACCGT (-4306 to -4292) |
| 468 | ATF4 | CTGCTGAGATTGCAGTA (-4933 to -4917)<br>ATCTTGAGAGAGCTCAT (-4449 to -4433)<br>ACCATGACTGGGCAAGC (-3612 to -3596)<br>TTGCTGACTGTGCTCCC (-3105 to -3089)<br>GGACTGACTTGGCTGAG (-2940 to -2924)<br>ATTTGCACAGTCATCTG (-2230 to -2214)<br>CCTCTGAGGCAGCAGGA (-1788 to -1773)<br>CCATGCAGACTCAGCCG (-893 to -877) | GGCGTGAGTCAAGGG (+513 to +527) |

**Note: TSS, transcriptional start signal and TIS, translation initiation signal.**

### Table S2. The primer pairs used for qRT-PCR analysis

| Name | Forward primer (5' to 3') | Reverse primer (5' to 3') |
|---|---|---|
| β-actin | CATGTACGTTGCTATCCAGGC | CTCCTTAATGTCACGCACGAT |
| Nrf1 | GCTGGACACCATCCTGAATC | CCTTCTGCTTCATCTGTCGC |
| Nrf2 | TCAGCGACGGAAAGAGTATGA | CCACTGGTTTCTGACTGGATGT |
| GCLM | GTGTGATGCCACCAGATTTGAC | CACAATGACCGAATACCGCAGT |
| HO-1 | CAGAGCCTGGAAGACACCCTAA | AAACCACCCCAACCCTGCTAT |
| Chop | GGAAACAGAGTGGTCATTCCC | CTGCTTGAGCCGTTCATTCTC |
| Bip | GAACGTCTGATTGGCGATGC | ACCACCTTGAACGGCAAGAA |
| ATF6 | AGCAGCACCCAAGACTCAAAC | GCATAAGCGTTGGTACTGTCTGA |
| ATF4 | CCCTTCACCTTCTTACAACCTC | TGCCCAGCTCTAAACTAAAGGA |
| XBP1 | CCCTCCAGAACATCTCCCCAT | ACATGACTGGGTCCAAGTTGT |
| IRE1 | GAGACCCTGCGCTATCTGAC | CTTGGCCTCTGTCTCCTTGG |
| PERK | CTTCCAGTGGGACCAAGACC | CGAGGTCCGACAGCTCTAAC |
| PSMB6 | TCAAGAAGGAGGGCAGGTGT | GTAAAGTGGCAACGGCGAA |
| PSMA1 | ATTCATCAAATTGAATATGCAAT | CTCTGATTGCGCCCTTTTCAA |
| PSMA4 | TTGCTGTACATTGGCTGGGA | ACACAGCTGCAGCGCTATTA |
| PSMA7 | TACATCACCCGCTACATCGC | AGAGCCTAGGAGTGCCATCA |
| PSMB7 | CTGTGTCGGTGTATGCTCCA | TGCCAGTTTTCCGGACCTTT |
| PSMC1 | ACAAGGTGCATGCCGTGATA | CTGTGCCAGGTGGACCATAG |
